# Probability waves: pattern-based p-value correction in mass univariate analysis between two event-related potential waves

**DOI:** 10.1101/2019.12.12.873570

**Authors:** Dimitri Marques Abramov

**Author notes:** Address: Avenida Rui Barbosa, 716. Bairro Flamengo, Rio de Janeiro, RJ, CEP 20021-140.

## Abstract

**Background:** Methods for p-value correction are criticized for either increasing Type II error or improperly reducing Type I error. This problem is worse when dealing with hundreds or thousands of paired comparisons between waves or images which are performed point-to-point. This text considers patterns in probability vectors resulting from multiple point-to-point comparisons between two ERP waves (mass univariate analysis) to correct p-values. These patterns (probability waves) mirror ERP waveshapes and might be indicators of consistency in statistical differences.

**New method:** In order to compute and analyze these patterns, we convoluted the decimal logarithm of the probability vector (p’) using a Gaussian vector with size compatible to the ERP periods observed. For verify consistency of this method, we also calculated mean amplitudes of late ERPs from Pz (P300 wave) and O1 electrodes in two samples, respectively of typical and ADHD subjects.

**Results:** the present method reduces the range of p’-values that did not show covariance with neighbors (that is, that are likely random differences, type I errors), while preserving the amplitude of probability waves, in accordance to difference between respective mean amplitudes.

**Comparison with existing methods:** the positive-FDR resulted in a different profile of corrected p-values, which is not consistent with expected results or differences between mean amplitudes of the analyzed ERPs.

**Conclusion:** the present new method seems to be biological and statistically more suitable to correct p-values in mass univariate analysis of ERP waves.

## Introduction

When we analyze event-related potentials (ERP), we primarily focus on latencies and amplitudes of the arbitrarily determined elements in these waves. We generally calculate the maximum and mean amplitudes (mean of amplitudes within an interval) in these elements, and statistically infer the difference in these parameters between two samples of that wave. Wave latency is determined based on its maximum amplitude. These elements are the components (or waves) of a set of signals that are systematically observed in a population of individuals under the same experimental conditions. For instance, the P100 wave obtained from human brain activity under visual stimulation of a reverse pattern [1]. However, the definition and delimitation of this ERP is historically arbitrary “to the naked eye”. Moreover, traditional p-correction analyses have lost several pieces of information regarding the identity of these waves, e.g., differences between periods and phases of these ERPs.

Mass univariate analysis (MUA) brings a new perspective to assess ERP behavior, as it consists of describing the differences between two waves that are explored when they are compared point to point, and we plot the statistical differences, using a method we call here raster pairwise comparison (RPC), which returns a raster diagram of p-values as a function of time [2]. See figure 1. Hence, by using RPC we can observe the difference between waves over time, combining latencies and amplitudes in one single measurement [3].

**Figure 1.**
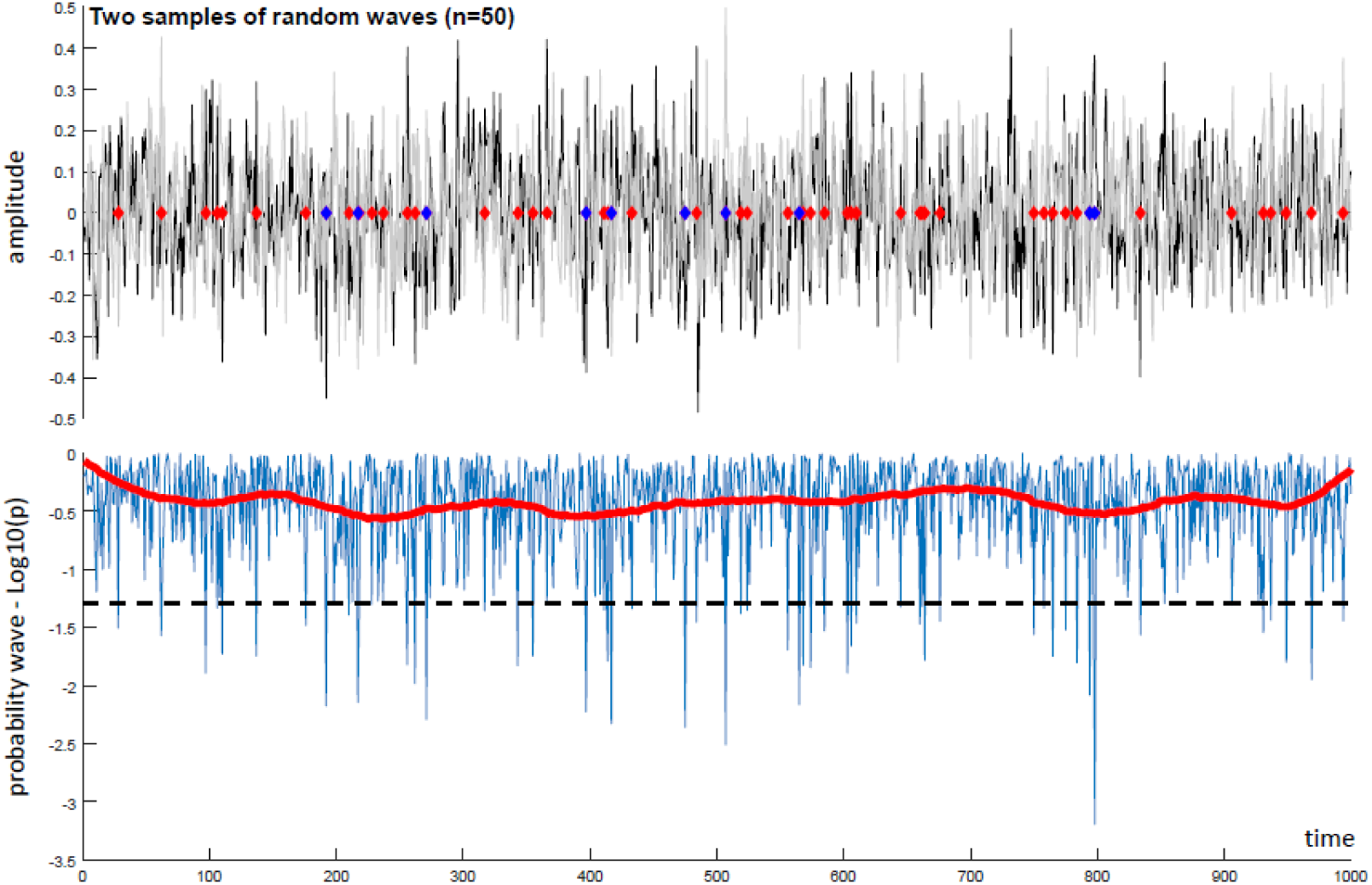
Mass univariate analysis with and without p-correction in a random scenario. Top: raster diagram of multiple pairwise comparisons (t-tests) between two samples with 50 random waves each (means in black and grey), and 1000 time points. Each blue diamond represents p ≤ 0.01, while the red ones represent p ≤ 0.05 regarding the null hypothesis for each pair of point vectors. We found 57 significant differences (p < 0.05, ∼ 5% of all points). Bottom: non-corrected probability vector (blue) and corrected one (red) by the present method, observing that all significances were rejected.

These pairwise comparisons can be tested with the most suitable statistical method according to sample features. In MUA, hundreds, even thousands of comparisons are performed according to the temporal extensions of waves. These comparisons are explanatory, i.e., we broadly seek for differences. This brings us to the issue of multiple comparisons, which substantially increase type I errors [4]. Historically, the concern with spurious differences has led to the development of methods to correct the probability of having differences due to the number of comparisons. The most traditional method is Bonferroni correction. However, as other methods of the same class, it is quite conservative [4].

Benjamini & Hockberg developed a method of False Discovery Rate (FDR), which is less conservative and is indicated, along with its variations, for multiple comparisons of MUA scale [5, 6]. After, other variants of FDR method were developed, which concern interdependence among comparisons [7, 8]. These correction methods are used to compare, e.g., genomes and resonance imaging (pixel to pixel). We used this method in a previous study for a point-to-point comparison of ERP waves (RPC).

Although FDR methods presume that pairwise compared data vectors are correlated to each other in terms of covariance (which would explain the fact that they correct p-values with a lower degree of rigor), these methods do not consider covariance of sampling vectors when calculating p-value correction.

We propose here a new paradigm for the correction of multiple comparisons in test sets where the variables tested have a natural correlation to each other, which can be considered *a priori*. P-values form a probability vector with behavioral patterns that might indicate that differences are statistically consistent in wave regions with p ≤ α.

### Theory and Method Application

A probability vector might have a visibly stochastic profile (figure 2), in which the distribution of statistically significant p-values does not follow any pattern. In order to observe this behavior, we derived the original p-value vector into a *p*’ vector, where *p*’ *= log*_*10*_*(p).* This vector, as a whole, suggests that these significant p-values might be erratic (i.e., type I errors).

**Figure 2.**
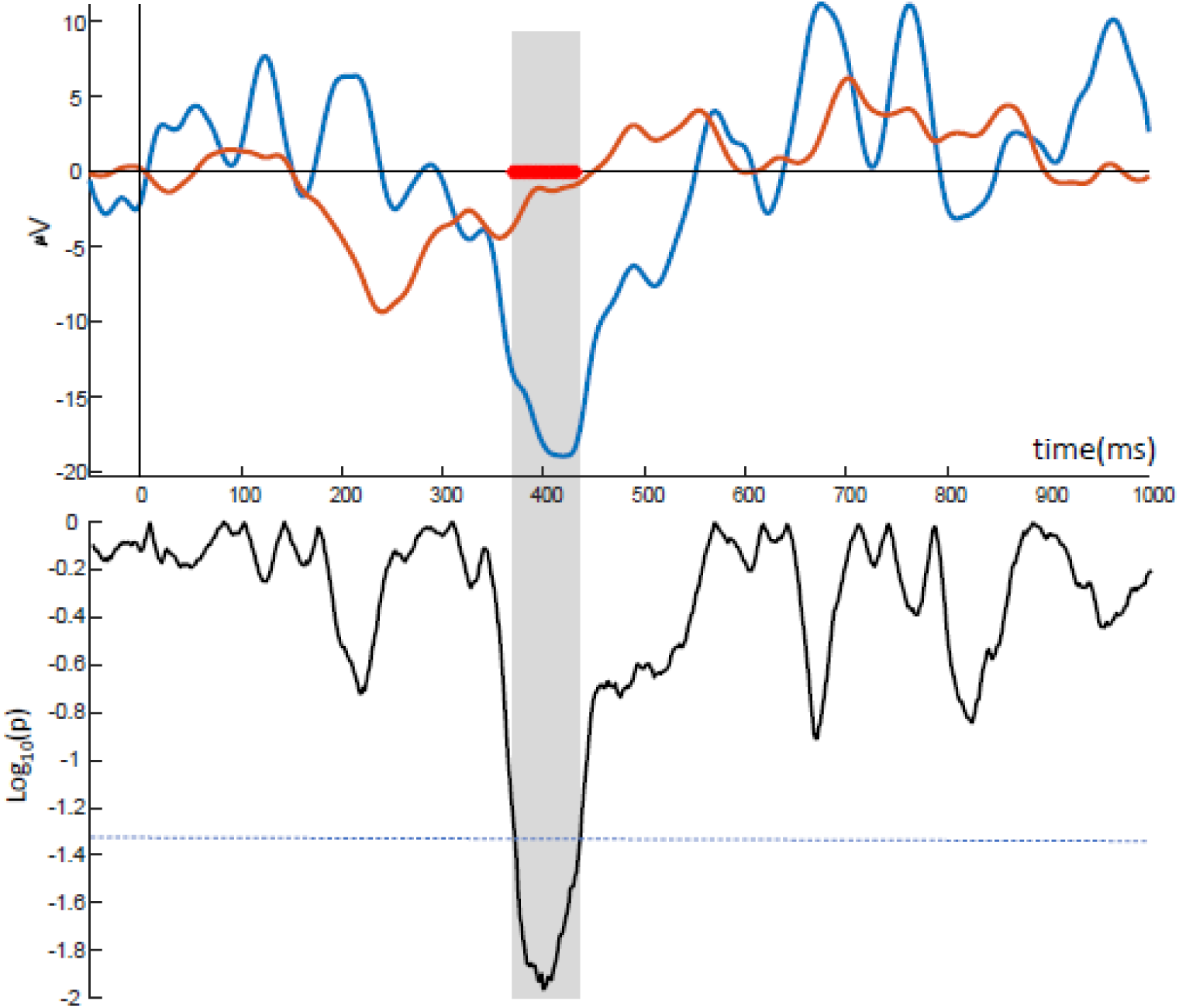
Probability waves. Vector of probability of rejecting the null hypothesis (bottom), from the raster pairwise comparison between two ERPs (top), showing collective behavior patterns of p’-values, organizing “probability waves” with profiles similar to the ERP. Shaded areas are the ones where the null hypotheses were rejected by α = 0.05 (red diamonds). p’ = log_10_(p).

However, another probability vector might show patterns of order that denote covariance between sampling data of neighbors (figure 3). This pattern observed is the gradual and massive evolution of *p*’-values forming a “probability wave” equivalent to that of the analyzed ERP (figure 3, dashed window).

**Figure 3.**
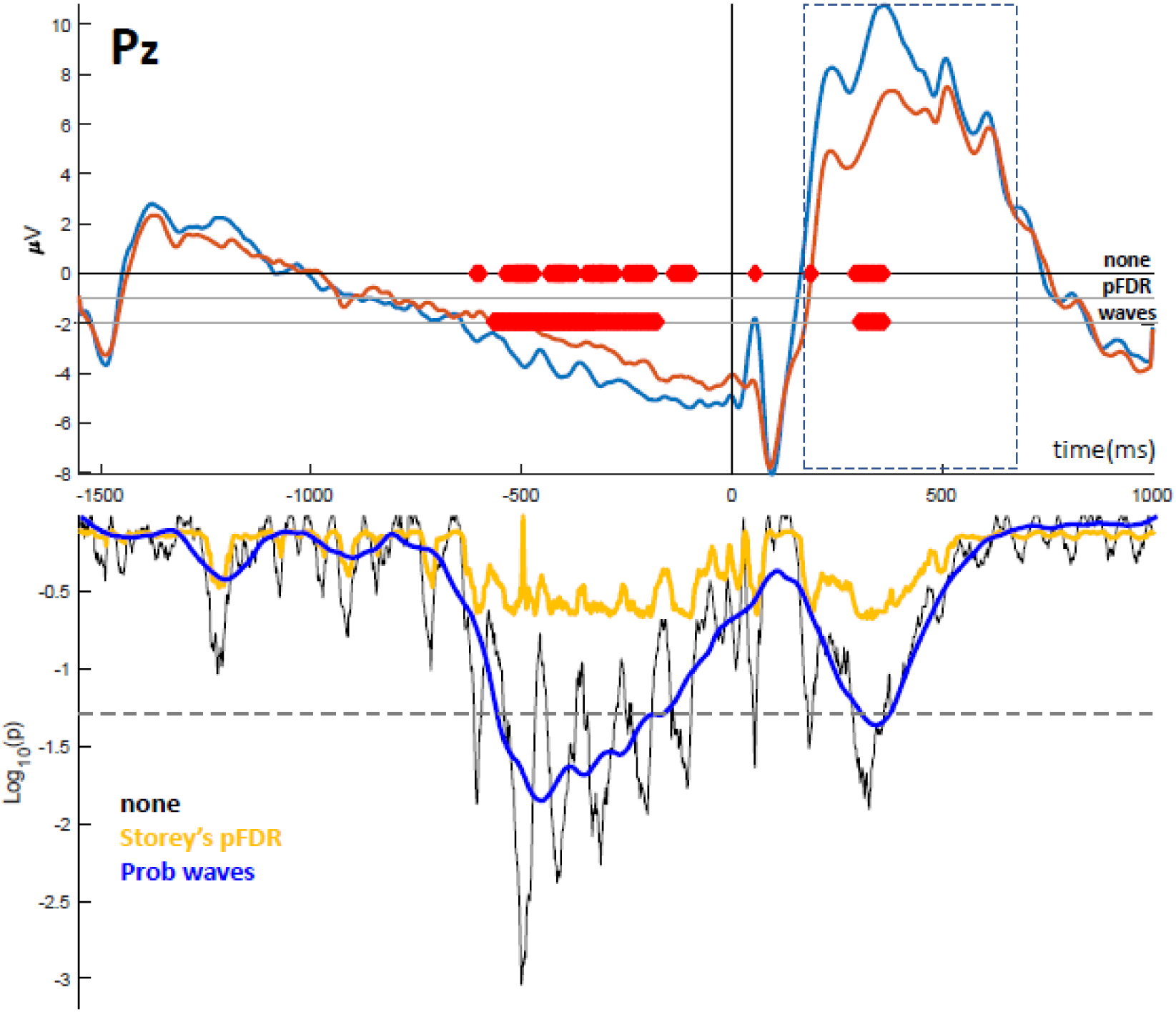
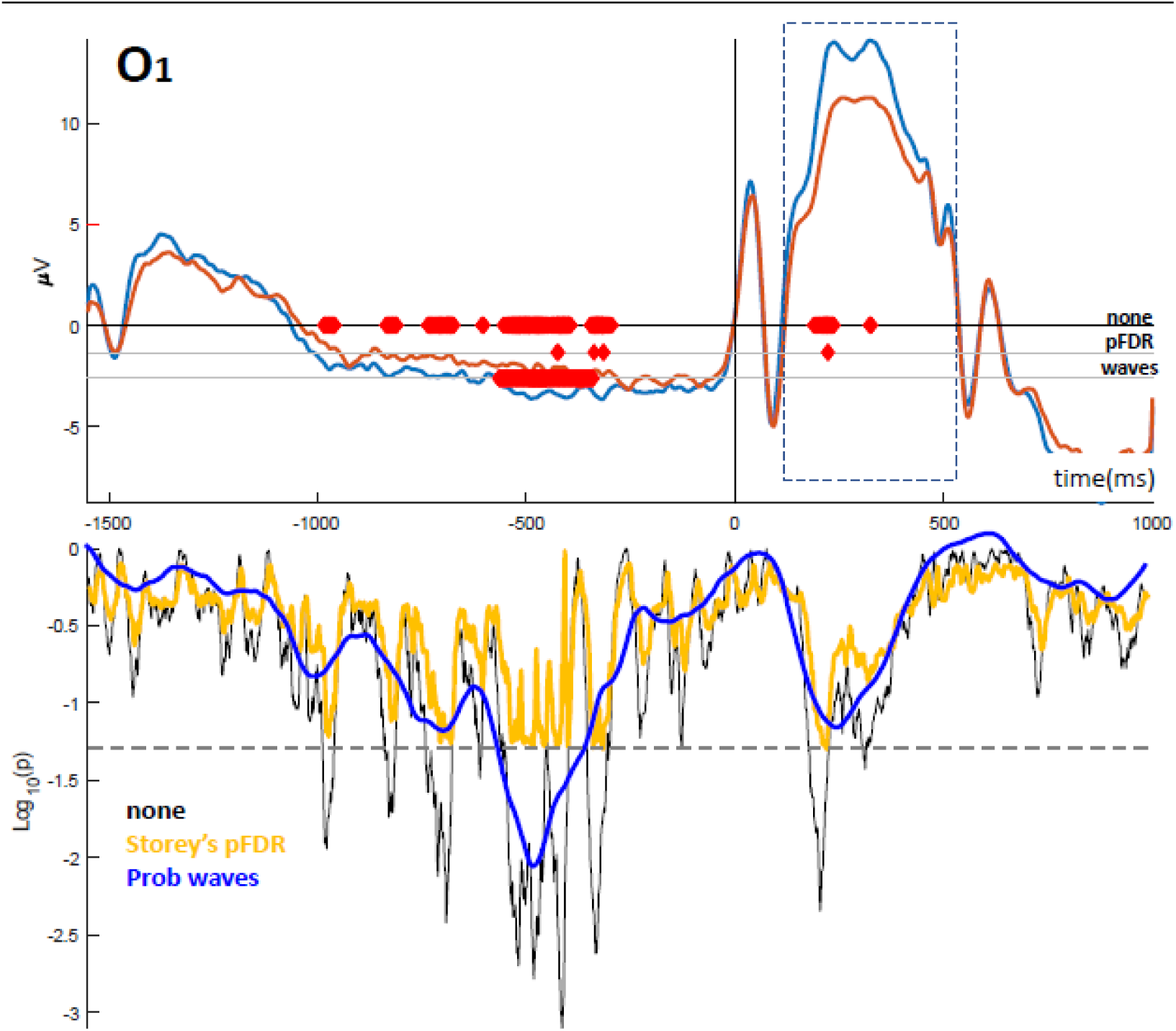
Methods for p-correction in UVA. Top: comparing waves from Pz electrode between groups (control in blue, cases in red), correlated to Attention Network Test behavior, performed by typical and ADHD youths (see text). P3 wave is inside the dashed window. The non-corrected probability wave (bottom, black) shows several rejected null hypotheses (red diamonds: p < 0.05). P-correction was performed by positive FDR (“pFDR”, orange) and the proposed method (“wave”, blue). The gray dashed line corresponds to the statistical boundary (p < 0.05). Bottom: for waves from O1 electrode.

We are not analyzing these behavioral patterns with unaided eye. One way of isolating such patterns is using mathematical convolution of *p*’-value in the *t*_*o*_*-t* interval, using a vector of n-values, according to the equation:

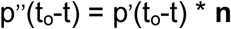

where *t*_*o*_ = [1, 2,…*t*_*max*_*-t*], *t*_*max*_ is the total size of the wave under analysis (in time points) and **n** is a value vector that sets one regular curve with size *t*:

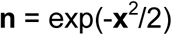

where **x** is the vector [*x*_*1*_,*x*_*2*_,…,*x*_*t*_], where *x*_*1*_ = -√2, *x*_*t*_ = √2, and *t* is the number of points in the wave compatible with the period of ERPs to be compared. We used here 60 points (equal to 100ms, at a 600Hz sampling rate). In the convolution algorithm, **n** slides on the **p**’ vector, thus creating a smoothing wave, where erratic values (with periods much lower than t) tend to be suppressed. We studied the effect of the convolution of **p’** by **n** (*t* = 60 bins ∼ 100ms) on the probability waveform resulting from multiple U-tests (Mann-Whitney) between two ERP waves (case and control), from the O1 and Pz electrodes (10-20 montage), obtained in an experiment that evaluated neurophysiological correlates of behavior in the Attention Network Test of both typical (n = 20) and ADHD children (n = 19) [3]. We show the uncorrected probability waves (figure 3, bottom of Pz and O1 panels, in black) and those corrected by the proposed method (blue). We also used the Storey method for p-correction (positive False Discovery Rate, p-FDR, orange) [9], which is less conservative than the Benjamini-Hockberg [6] and Benjamini-Yekutieli [7,8] methods.

To convolve over the probability wave, we used a normal curve with a period of *t* = 100ms, with order of magnitude of the expected waves and that is thus able to isolate patterns with periods equal to or greater than 100ms. To estimate the reliability of p-correction methods, we compared the mean amplitude of the cognitive target-related potentials (dashed line) on the Pz electrode, which is the P300 wave [10], and on the O1 electrode. We tested the null hypothesis between those mean amplitudes using the Mann-Whitney U Test.

Observing the effect of the proposed method on a stochastic scenario (figure 1), all statistical differences were rejected, as they should be. After convolution, the resulting probability wave shows very low amplitudes (figure 1, bottom, red wave).

In the real-life scenario, the mean amplitudes of the target-related potential from Pz were significantly different between groups (control: 6.70 ± 3.16 μV; cases: 4.18 ± 4.24 μV; p = 0.028). However, significance was not observed in the corresponding differences from O1 electrode (control: 9.66 ± 2.52 μV; cases: 7.82 ± 3.17 μV; p = 0.053). As observed in figure 3, the p-correction methods resulted in different **p’**-vectors. Here, the proposed method rejected the null hypotheses, thus behaving consistently with the differences found between the mean amplitudes.

## Discussion

The P3 wave is an attention-related cognitive neurofunctional complex that manifests on parietal site, and which appears around 300 milliseconds after the corresponding event [10]. Thus, we would expect some effect of the test on the P3 wave only on the Pz electrode. The proposed method corrected the p-values more consistently with the expected behavior of the P3 wave than the pFDR method. In the universe of event-related potentials, the method of probability wave using convolution of compatible magnitude to the biologically expected one was more reliable.

We presume that in Nature, collateral points of a biological wave are correlated to each other, as they derive from deterministic processes, although they have a chaotic nature. Thus, the behaviors of both equivalent waves, which are produced by the same source, follow the same causal mechanisms. The differences between these waves, statistically speaking, would also follow, by principle, a variation pattern that mirrors the profile of these waves.

Considering waves resulting from two different processes derived from the same causal mechanism (for instance, potential related to rare and frequent stimuli in an OddBall paradigm, resulting from different neural processes derived from the same neural mechanism [10]), theoretically, the chance of observing a false negative test result (type II error) is much lower than a false positive result (type I error). This is because different processes have a causal relationship with the same mechanisms (non-randomization).

Therefore, since a set of points is statistically different (rejecting the null hypothesis), because they correspond to the lowest values in a subset of points of the probability vector, which shows an organized behavioral pattern (a probabilistic wave), these statistically determined differences might be considered to be true.

Hence, in a Mass Univariate Analysis between two ERP waves presumably derived from the same biological processes, values lower than log10(α) of the probability pattern vector (p’’) do not correspond to type I errors.

## Acknowledgements

I would like to especially thank prof. Mario Fiorani Jr, whose computational function for convolution was used in this work. I would also like to thank the oswaldo cruz foundation for the opportunity to develop this work.

The author authors have no competing interests to declare.

